# Liquid-liquid phase separation of florigen activation complex induces flowering

**DOI:** 10.1101/2021.08.19.456992

**Authors:** Ken-ichiro Taoka, Hiroyuki Tsuji, Suai Anzawa, Mayu Enomoto, Yuka Koizumi, Juri Nakamura, Mari Tanaka, Akiko Fujita, Kyoko Furuita, Takashi S. Kodama, Masaru Fujimoto, Kazuo Kurokawa, Takashi Okamoto, Toshimichi Fujiwara, Akihiko Nakano, Chojiro Kojima

**Affiliations:** Kihara Institute for Biological Research, Yokohama City University, Yokohama, Japan; Graduate School of Engineering Science, Yokohama National University, Yokohama, Japan; Graduate School of Biological Sciences, Nara Institute of Science and Technology, Nara, Japan; Institute for Protein Research, Osaka University, Suita, Osaka, Japan; Graduate School of Agricultural and Life Sciences, The University of Tokyo, Tokyo, Japan; Live Cell Super-Resolution Imaging Research Team, RIKEN Center for Advanced Photonics, Wako, Saitama, Japan; Department of Biological Sciences, Tokyo Metropolitan University, Minami-osawa, Hachioji, Tokyo, Japan

## Abstract

Floral transition, regulated by the systemic action of the mobile florigen protein FLOWERING LOCUS T (FT), is essential for successful plant reproduction^1^. How FT controls downstream gene expression remains incompletely understood, although it relies on the florigen activation complex (FAC), a core component of FT function^2–4^. The FAC is a nucleus-localized transcriptional activator of genes encoding MADS-box transcription factors critical to reproductive development and consists of florigen FT; a scaffold 14-3-3 protein that is a key component for complex assembly; and FD, a basic leucine-zipper protein that recruits the FAC to DNA. Here we report that the FAC exhibits phase separation. In rice shoot apical meristem cells, rice florigen Heading date 3a (Hd3a) fused to the green fluorescent protein formed speckles in the nucleus. The FAC speckle is formed in a FAC-dependent manner in tobacco cells. Recombinant Hd3a, but not OsFD1, phase-separated in vitro, and this effect was enhanced in the presence of 14-3-3 protein. Furthermore, mutations affecting functionally important residues in the pocket region or C-terminal disordered region of Hd3a affected FAC phase separation, providing a biochemical framework for the protein’s effect on flowering. The ability to form condensates via phase transition represents a previously unknown mechanism for gene activation by the FAC.

## Main

Florigen is a systemic signal that induces flowering^5,6^ whose molecular identity was elucidated with the cloning of Arabidopsis (*Arabidopsis thaliana*) *FLOWERING LOCUS T* (*FT*) and orthologues from other plant species, such as *HEADING DATE 3A* (*Hd3a*) in rice (*Oryza sativa*)^1,7,8^. *FT, Hd3a* and orthologues are expressed and translated in leaves and then transported to the shoot apical meristem (SAM) responsible for the production of all above-ground tissue^7,9,10^. Next, the SAM is converted from a leaf-forming vegetative meristem to a flower-forming reproductive meristem through the formation of the florigen activation complex (FAC), comprising the florigen itself, the florigen receptor 14-3-3 protein and the basic leucine zipper (bZIP) transcription factor FD^2–4,11,12^. Although great strides have been made in understanding florigen function, the biochemical mechanisms by which the FAC activates downstream gene expression are not well understood. As a first step toward elucidating these, we determined the subcellular localization of the FAC. We performed super-resolution imaging microscopy by stimulated emission depletion (STED)^13,14^ of cells in the rice SAM. Because Hd3a is expressed in the phloem of vascular bundles^7,15,16^, we generated rice plants expressing *green fluorescent protein* (*GFP*) or *Hd3a-GFP* driven by the *rolC* promoter from *Agrobacterium rhizogenes*, which is highly active in the phloem ^7,17,18^. The resulting translated GFP and Hd3a-GFP were transported to the SAM, where they accumulated in cell nuclei (Fig. 1a,b). As a control for STED, we determined the localization of histone H2B fused to GFP (H2B-GFP)^19^ in the nucleus of SAM cells, revealing punctate signals and exclusion from the nucleolus indicative of the nucleosome (Fig. 1c). Free GFP showed a faint signal in the cytoplasm and was uniformly distributed in the nucleus, whereas Hd3a-GFP formed speckle-like structures in the nucleus (Fig. 1d,e). To confirm the formation of Hd3a-containing speckles, we visualized GFP fluorescence from the nuclei of SAM cells of transgenic plants harbouring the *Hd3apro:Hd3a-GFP* transgene^7^ by super-resolution confocal live imaging microscopy (SCLIM)^20,21^, a super-resolution imaging technique based on principles distinct from those underlying STED. However, as observed by STED, Hd3a-GFP defined a speckle-like structure in the nuclei of rice SAM cells (Fig. 1f,g,h).

**Fig. 1.**
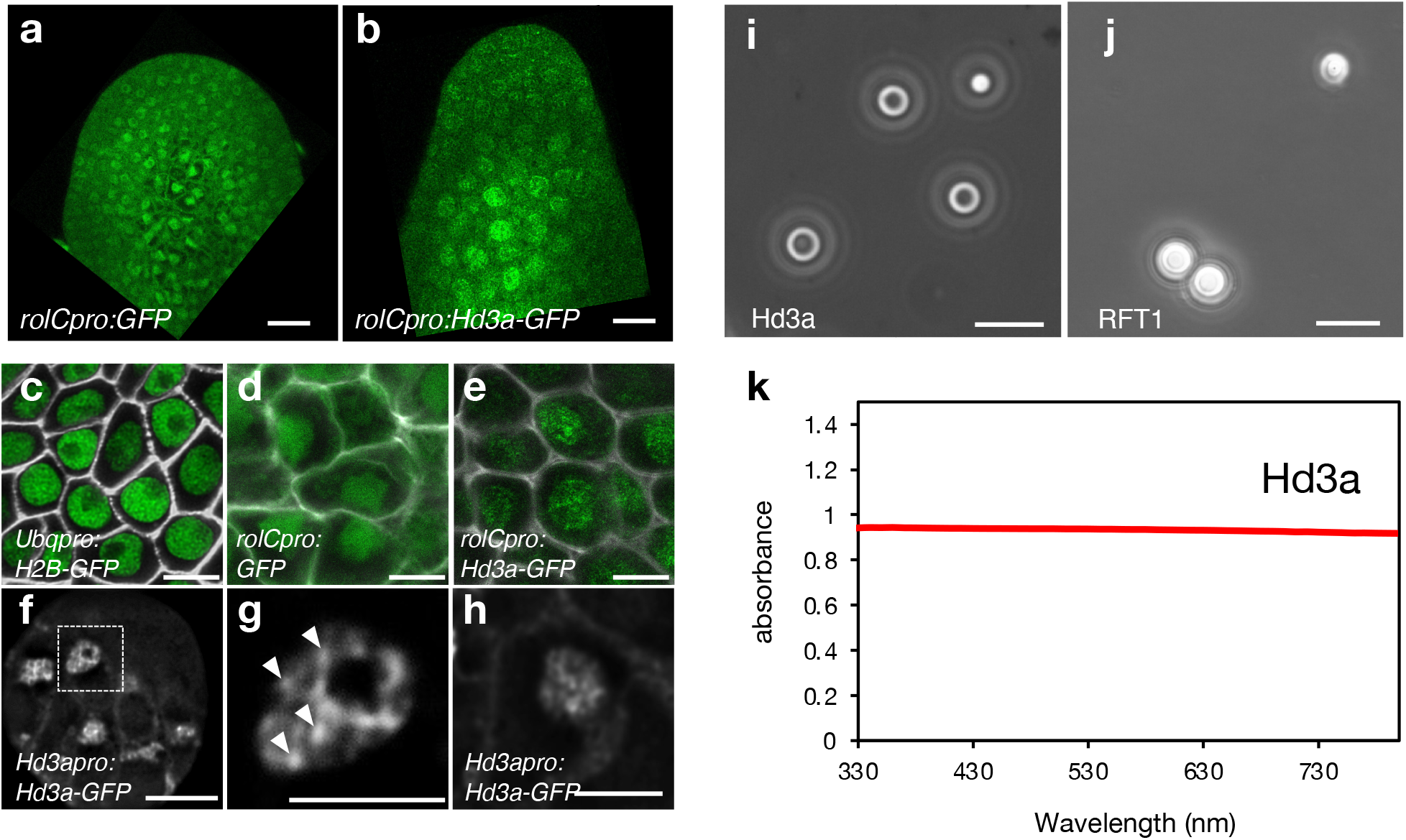
Hd3a forms nuclear speckles in the shoot apical meristem. **a**,**b**, Representative GFP fluorescence pattern from *GFP* (**a**) or *Hd3a-GFP* (**b**) constructs driven by the phloem-specific *rolC* promoter in the rice shoot apical meristem. Scale bars, 10 μm. **c–e**, Stimulated emission depletion (STED) imaging of cells within the rice shoot apical meristem harbouring the transgenes *Ubqpro:H2B-GFP* (**c**), *rolCpro:GFP* (**d**) and *rolCpro:Hd3a-GFP* (**e**). **f**, Super-resolution confocal live imaging microscopy (SCLIM) imaging of cells in the rice shoot apical meristem harbouring the transgene *Hd3apro:Hd3a-GFP*. Scale bars, 5 μm. **g**, Enlarged view of the region highlighted by a dashed box in (**f**). **h**, Enlarged view of a nucleus in the shoot apical meristem from a *Hd3apro:Hd3a-GFP* transgenic rice line. **i**,**j**, Spherical condensates of Hd3a (**i**) and RFT1. Scale bars, 10 μm. (**j**). **k**, Absorbance of recombinant Hd3a in a PEG-containing solution.

Cellular speckle-like structures have recently been proposed to constitute biomolecular condensates, i.e., phase separation^22–27^. To assess whether the observed speckle-like Hd3a structures are caused by liquid condensates, we purified recombinant Hd3a and its paralogue rice FT1 (RFT1) and examined their ability to form condensates in vitro. Indeed, both recombinant Hd3a and RFT1 proteins appeared to form condensates, with a diameter of about 10 µm, in vitro in solutions containing polyethylene glycol (PEG) (Fig. 1i,j). To characterize these condensates, we measured their absorbance in the range of 330–800 nm, as absorbance over this range indicates the degree of condensation and reflects the size of the dispersed condensates in solution: larger condensates show a constant absorbance, as illustrated with glass beads of different diameters in solution (Extended Data Fig. 1). Hd3a showed a constant absorbance over the measurement range (Fig. 1k) that was reminiscent of the pattern seen for large-size glass beads, suggesting the presence of large condensates containing Hd3a and RFT1 (Extended Data Fig. 2). This result was consistent with microscopic observations of the larger condensates with roughly 10-µm condensates. Furthermore, recombinant Hd3a and RFT1 formed spherical condensates rather than aggregated precipitates (Fig. 1i,j), indicating that Hd3a and RFT1 undergo liquid phase separation in vitro.

Transient expression of *Hd3a* and *OsFD1* in rice protoplasts reconstituted the FAC by complexing with endogenous 14-3-3 proteins, leading to the activation of one of its transcriptional targets, *OsMADS15*, which encodes a MADS-box transcription factor^2,28^ (Fig. 2a). FAC reconstitution has also been reported from the transient expression of Arabidopsis *FT* and *FD* or rice *FD4* and *RFT1* in *N. benthamiana* leaves^11,29^. To investigate whether nuclear speckles form upon reconstitution of the FAC, we transiently expressed *Hd3a-GFP* and *OsFD1* in *N. benthamiana* leaves and determined the green fluorescence pattern detectable in their nuclei. The mean number of speckles per nucleus was small (less than 0.5) when *Hd3a* alone or *OsFD1* alone was expressed, while nuclei contained an average of three speckles upon co-expression of *Hd3a* and *OsFD1* (Fig. 2b,c). This observation suggests that FAC formation entails the localization of Hd3a into nuclear speckles.

**Fig. 2.**
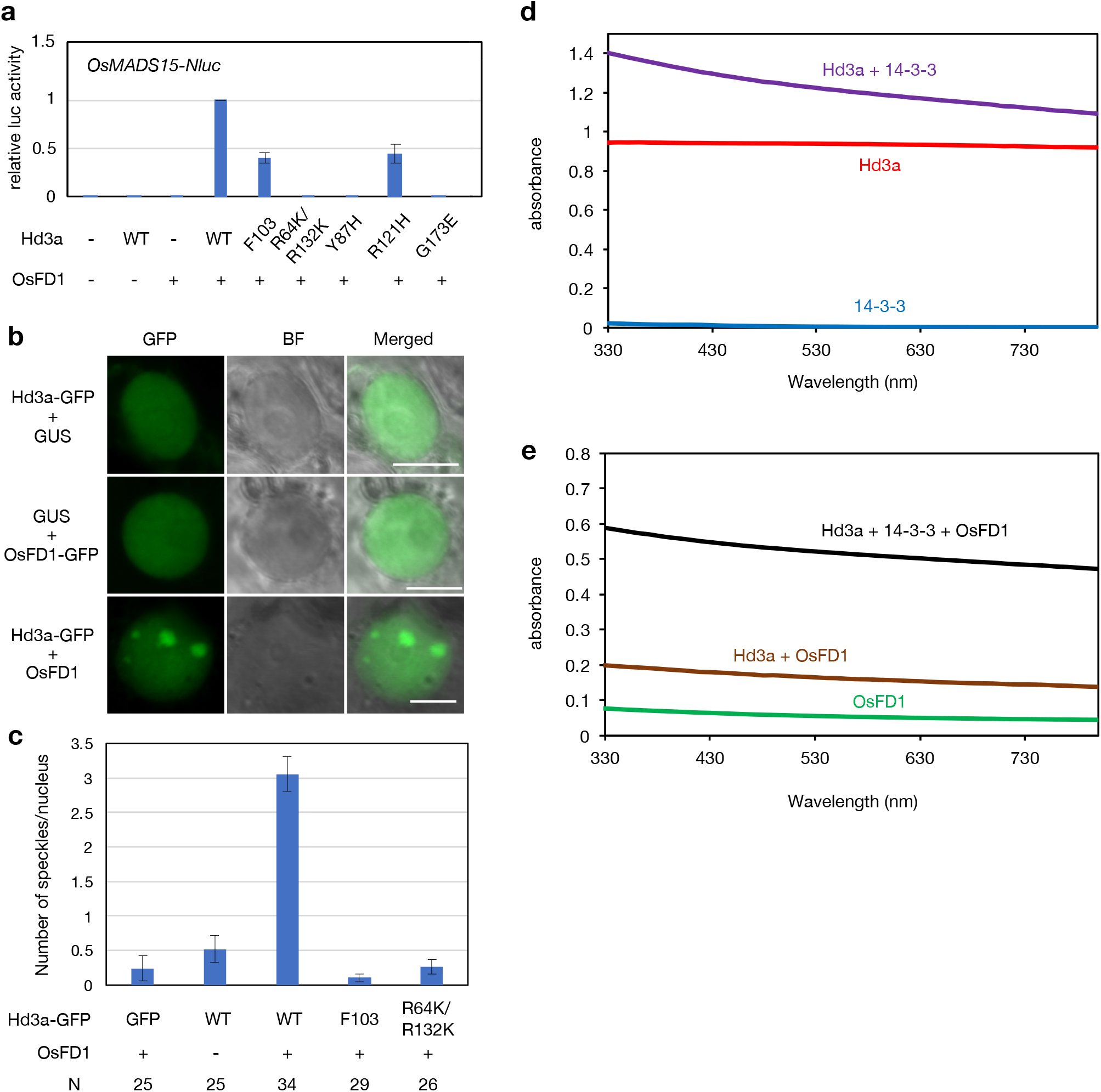
Formation of the florigen activation complex (FAC) facilitates nuclear speckle formation and in vitro condensation. **a**, Effects of Hd3a mutations on *OsMADS15* transcriptional activation. **b**, Hd3a-GFP localization in the nucleus of *N. benthamiana* leaf epidermal cells. Scale bars, 10 μm. **c**, Effects of Hd3a mutations on the mean number of nuclear speckles. **d**,**e**, Absorbance of recombinant Hd3a alone (red), GF14c alone (blue) and Hd3a with GF14c (purple) (**d**) and of OsFD1 alone (light green), Hd3a with OsFD1 (brown) and Hd3a with OsFD1 and GF14c (black) (**e**).

To characterize whether these nuclear speckles formation is explained by condensate formation in vitro, we measured the absorbance of recombinant Hd3a, 14-3-3 protein GF14c^2^ and OsFD1 alone or in mixtures over the range of 330–800 nm. We were unable to prepare a large amount of full-length OsFD1, but we succeeded in preparing a C-terminal truncated version of OsFD1 that retains the coiled-coil region. Hd3a showed constant absorbance over the measurement range, whereas 14-3-3 and OsFD1 showed little absorbance on their own (Fig. 2d,e). To determine the effect of 14-3-3 protein on condensate formation by Hd3a or OsFD1, we added 14-3-3 protein to recombinant Hd3a or OsFD1 and measured the absorbance of the resulting mixtures. The absorbance was higher when 14-3-3 was included than with recombinant Hd3a or OsFD1 protein alone (Fig. 2d,e). We next added recombinant 14-3-3 protein to a mixture of recombinant Hd3a and OsFD1 proteins and measured the absorbance of the resulting solution (Fig. 2e), finding that the mixture displayed a constant absorbance of about 0.2. The absorbance increased considerably when recombinant 14-3-3 protein was added to the mixture (Fig. 2e). Increasing ratio of Hd3a against 14-3-3 resulted in higher absorbance (Extended Data Fig. 3). These results suggest that the formation of the three-protein FAC promotes the appearance of condensates in vitro.

To better understand FAC function, we examined whether the interaction between Hd3a and 14-3-3 is required to form nuclear speckles. Accordingly, we introduced mutations in Hd3a that abolish its interaction with 14-3-3 proteins. The phenylalanine 103 (F103), arginine 64 (R64) and arginine 132 (R132) residues of Hd3a form a hydrogen bond that is essential for interaction with 14-3-3 proteins, and their mutation to create Hd3a^F103A^ or Hd3a^R64K,R132K^ mutants abolished the interaction between Hd3a and 14-3-3 interaction and resulted in a loss of FAC transcriptional activity^2^ (Fig. 2a). Neither *Hd3a*^*F103A*^*-GFP* nor *Hd3a*^*R64K,R132K*^*-GFP* led to the formation of nuclear speckles when transiently expressed in *N. benthamiana* leaves (Fig. 2c, 3a). To investigate whether the interaction between Hd3a and 14-3-3 is necessary for condensate formation, we performed an in vitro assay in which we added recombinant 14-3-3 protein to mutant recombinant Hd3a and measured the absorbance of the solutions. The absorbance of recombinant Hd3a^F103A^ and Hd3a^R64K,R132K^ in the presence of recombinant 14-3-3 protein was lower than that of intact Hd3a with 14-3-3 (Fig. 3b,c). These results indicate that the interaction between Hd3a and 14-3-3 is a prerequisite for condensate formation.

**Fig. 3.**
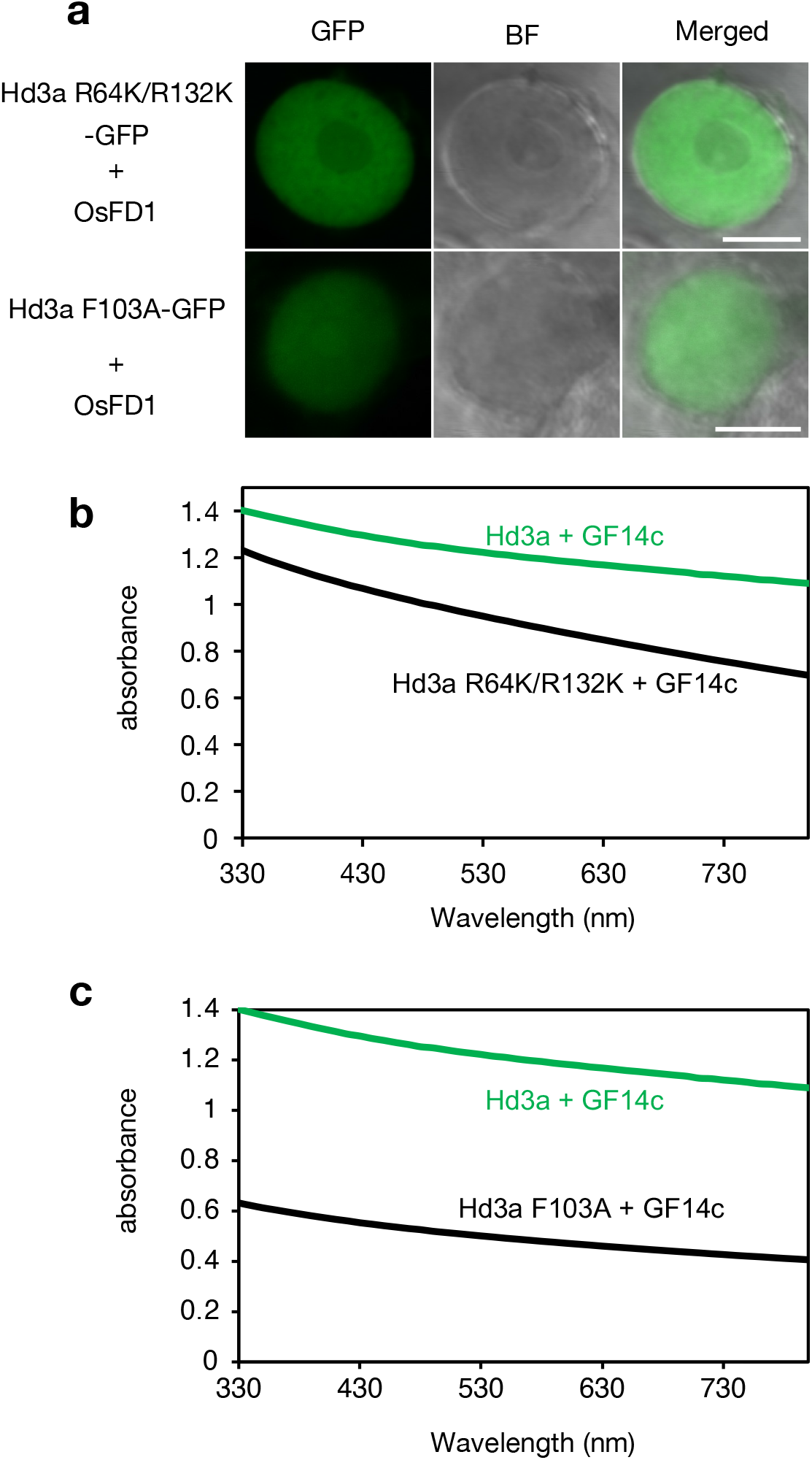
Interaction with 14-3-3 protein facilitates nuclear speckle formation and in vitro condensation of Hd3a. **a**, Localization pattern of Hd3a^R64K,R132K^-GFP and Hd3a^F103A^-GFP when co-expressed with *OsFD1* in the nucleus of *N. benthamiana* leaf cells. Scale bars, 10 μm. **b**,**c**, Absorbance of recombinant Hd3a with GF14c (green) and Hd3a^R64K,R132K^ with GF14c (black) (**b**) and of Hd3a with GF14c (green) and Hd3a^F103^ with GF14c (black) (**c**). Scale bars, 10 μm.

The family of phosphatidylethanolamine-binding proteins (PEBP), to which florigen belongs, is characterized by an anion-binding pocket that has been suggested to play an important role in the function of florigen^30,31^. The tyrosine 87 (Y87) residue is located deep in this potential ligand-binding pocket on the surface of florigen and is conserved across all FT orthologues, but is replaced by histidine in TERMINAL FLOWER 1 (TFL1) and its orthologues, floral repressors that are closely related to FT^32^. To explore the potential link between the pocket and condensate formation, we characterized a mutant version of Hd3a harbouring histidine instead of tyrosine at residue 87 (Y87H). Hd3a^Y87H^ failed to activate *OsMADS15* expression (Fig. 2a) but retained the ability to form nuclear speckles (Fig. 4a,b) and condensates in vitro (Fig. 4c). These results indicate that the Y87H mutation in the pocket region causes loss of transcriptional activation by the FAC without affecting nuclear condensate formation.

**Fig. 4.**
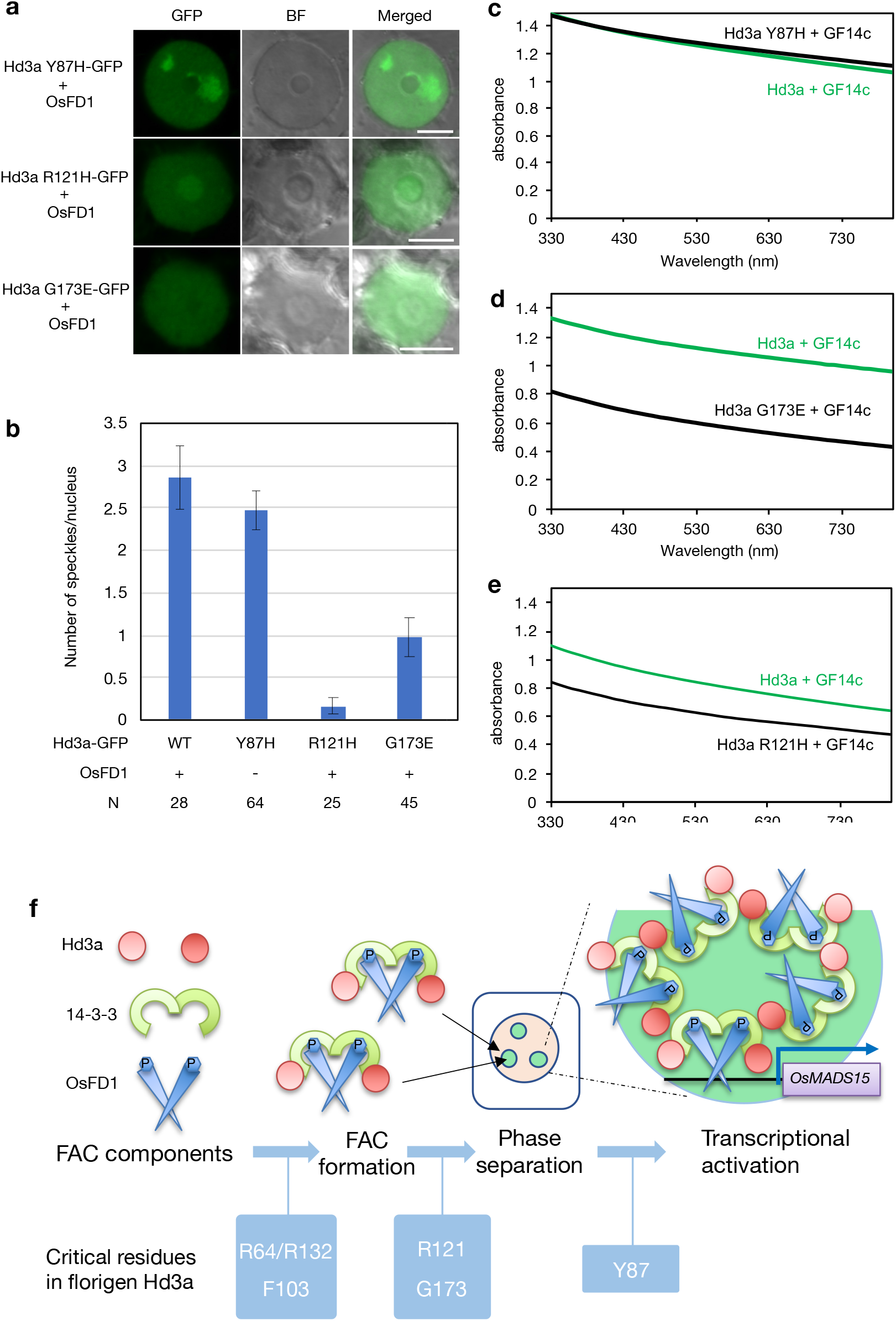
Substitutions in the potential ligand-binding pocket and C-terminal disordered region of Hd3a modulate nuclear speckle formation and in vitro condensation. **a**, Localization pattern of Hd3a^Y87H^-GFP, Hd3a^R121H^-GFP and Hd3a^G173E^-GFP when co-expressed with *OsFD1* in the nucleus of *N. benthamiana* leaf cells. Scale bars, 10 μm. **b**, Effects of Hd3a mutations on the number of nuclear speckles. **c**–**e**, Absorbance of recombinant Hd3a with GF14c (green throughout) and of Hd3a^Y87H^ with GF14c (black, **c**), Hd3a^R121H^ with GF14c (black, **d**) and Hd3a^G173E^ with GF14c (black, **e**), mixed 50:50 in each case. **f**, Proposed model of FAC activity through condensate formation by condensate formation in the nucleus. FAC formation induces condensate formation, which requires the interaction of Hd3a with 14-3-3 via the Hd3a residues R64, F103 and R132. G173 in the C-terminal disordered region and the internal R121 are essential for condensate formation after FAC formation. The potential anion-binding pocket of Hd3a includes Y87, which is essential for FAC function but is dispensable for condensate formation, suggesting that it acts downstream of condensate formation.

We also examined the link between FAC function and condensate formation for other known loss-of-function mutations, namely those corresponding to the Arabidopsis *ft-1* and *ft-3* alleles. The FT protein in the *ft-1* mutant contains two consecutive glycine residues (G) in the C-terminal disordered region, one of which is mutated to glutamic acid (E)^15,16,33,34^. A transient expression assay revealed that the equivalent mutant in rice Hd3a is G173E, as it failed to activate *OsMADS15* transcription (Fig. 2a). We next examined whether Hd3a^G173E^ can form nuclear speckles. We discovered that *Hd3a*^G173E^ could not form nuclear speckles when transiently expressed in *N. benthamiana* leaf cells (Fig. 4a,b). Likewise, recombinant Hd3a^G173E^ suppressed condensate formation when incubated with recombinant 14-3-3 protein in vitro (Fig. 4d). Since the G173 residue is located in the C-terminal disordered region of Hd3a^2,34^, this substitution does not affect the overall structure. Thus, the behaviour of the Hd3a^G173E^ mutant underscores the importance of the C-terminal disordered region following FAC formation.

The Arabidopsis *ft-3* allele is equivalent to a mutant of Hd3a in which arginine 121 is replaced by histidine (R121H)^15,16,33^. The R121 residue is located internally within the Hd3a structure. Hd3a^R121H^ can interact with 14-3-3 proteins^2^ but appeared to have partially lost its transcriptional activation potential (Fig. 2a). When we transiently expressed Hd3a^R121H^ in *N. benthamiana* leaf cells, we observed that it could not form nuclear speckles (Fig. 4a), and it inhibited condensate formation in the presence of 14-3-3 protein to a greater extent than intact Hd3a in vitro (Fig. 4e). Since R121 is embedded in the Hd3a molecule^2,34^, we investigated whether R121H disrupted the structure of Hd3a. Indeed, the structure of Hd3a^R121H^ appeared to be different from WT, since the nuclear magnetic resonance (NMR) spectrum of Hd3a^R121H^ was different from WT (Extended Data Fig. 4). These results suggest that R121H is a mutation that prevents condensate formation by changing the protein structure of Hd3a while maintaining its interaction surface with 14-3-3 proteins.

Collectively, our results suggest that the FAC exerts its transcriptional activation through a mechanism that involves nuclear speckles and condensation and that depends on the conserved R121 residue and the C-terminal disordered region of Hd3a (to which the G173 residue maps). Although the mechanism of condensate formation by florigen is currently unknown, we hypothesize that it involves a multivalent interaction such as charge interactions between FAC components. For example, the surface of florigen is basic, while that of 14-3-3 is acidic^2,35^. The electrostatic interactions based on these differences may induce a multivalent interaction between florigen and 14-3-3 to promote condensate formation. In addition, the phosphorylated SAP motif of FD may help bridge multiple 14-3-3 dimers to facilitate condensate formation through multivalent interactions.

The results of this study suggest that condensate formation serves as a mechanism by which the FAC activates downstream gene expression (Fig. 4f). Interaction between Hd3a and FD mediated by 14-3-3 protein is essential for condensate formation, and the Hd3a C-terminal disordered region containing the G173 residue promotes this process after FAC assembly. This multi-layered regulation along with condensate formation contributes to FAC-mediated transcriptional activation.

Phase-separated condensation of FAC may provide a cooperative switching mechanism for gene expression related to flowering. The phase-separated condensation of FAC occurs when all required proteins are simultaneously present in high concentrations in a cell: for FAC, Hd3a, 14-3-3 and OsFD1. In rice SAMs, 14-3-3 and OsFD1 accumulate in high concentrations in the cells during the vegetative phase^36^. Therefore, the phase-separated condensation of FAC may provide a cooperative switching mechanism whereby Hd3a is transported into the SAM and its concentration is increased, and the phase-separated condensation of FAC and transcriptional activation occur only when the concentrations of Hd3a, 14-3-3 and OsFD1 are sufficiently high.

## Methods

### Plant materials and growth conditions

Rice (*Oryza sativa* L. subspecies *japonica*) variety Nipponbare was used as wild type. *Hd3apro*:*Hd3a*–*GFP, rolCpro*:*Hd3a–GFP, rolCpro*:*GFP* and *Ubqpro*:*H2B–GFP* transgenic rice plants were described previously^7,19^. Transgenic rice cell lines derived from the *OsMADS15-nanoLuc* gene targeting line (*M15NL-KI*) were described previously^28^. Transgenic rice plants were generated using Agrobacterium (*Agrobacterium tumefaciens*)-mediated transformation of rice calli, as previously described, and hygromycin-resistant plants were regenerated from the transformed calli^7^. Plants were grown in growth chambers at 70% humidity, under short-day conditions with daily cycles of 10 h of light at 28 °C and 14 h of darkness at 25 °C. Light was provided by white fluorescent tubes (400–700 nm, 100 μmol m^−2^ ^s−1^). Cells from rice suspension cultures were maintained as described previously. *Nicotiana benthamiana* plants were grown under long-day conditions with daily cycles of 16 h of light at 22 °C and 8 h of darkness at 20 °C. Light was provided by white fluorescent tubes (400– 700 nm, 100 μmol m^−2^ s^−1^).

### Plasmid construction

To make an expression vector in rice protoplast, a 2005-bp fragment of maize ubiquitin promoter and a 252-bp fragment of NOS terminator were amplified by PCR. A 1784-bp of Gateway recombination site were obtained by digestion of pGWB26 vector^36^ with restriction enzymes, XbaI and SacI. These three fragments were inserted into pGreen II plasmid^37^ to make the expression vector, pGIIpUbqGWT7Ct. Mutations in Hd3a were introduced by PCR mutagenesis with a designed primer set. The resultant Hd3a mutants were cloned into pENTR-D-TOPO (Thermo Fisher Scientific) and introduced into pGIIpUbqGWT7Ct with LR clonase II (Thermo Fisher Scientific). For OsFD1 expression, pUbq-OsFD1^2^ was used. For Agroinfiltration in tobacco, pGWB602 or pGWB605^38^ were used for the destination vectors.

### Agroinfiltration

Agroinfiltration was performed as described previously^39^. The Agrobacterium strains EHA105 and MP90 were used for transformation of pGWB602 or pGWB605 derivatives^38^ and pBIC p19^40^, respectively. pBIC p19 harbours the silencing suppressor p19 from tomato bushy stunt virus (TBSV). After overnight growth in LB medium, agrobacteria were collected by centrifugation and resuspended to an OD_600_ = ∼0.6 in infiltration buffer (2 mg ml^™1^ MgCl_2_·6H_2_O, 150 µM acetosyringone, 10 mg ml^™1^ MES-KOH, pH 5.6). The appropriate combinations of cell suspensions were mixed in equal volumes and infiltrated into *N. benthamiana* leaves 4–5 weeks after germination with a 1-mL syringe. The cells were fixed with 4 % paraformaldehyde for 5-6 days after infiltration and observed within a month.

### Imaging by confocal laser scanning microscopy

The shoot apical meristems (SAMs) from transgenic rice plants were dissected under a microscope^41^. For imaging by stimulated emission depletion (STED), SAMs were fixed by 4% paraformaldehyde and cleared with ClearSee, stained by SCRI SR2200 for cell wall staining and visualized using a confocal laser-scanning microscope (TCS SP8; Leica Microsystems, Tokyo, Japan) equipped with 592-nm STED laser, a 405-nm laser and a pulsed white-light laser (WLL) line and 93· oil-immersion objective lens. Samples were excited with the 488-nm wavelength of the WLL (80 MHz) and depleted with the 592-nm STED laser. Collected images were deconvolved using the default settings of the STED module in Huygens Professional Deconvolution software. For imaging by super-resolution confocal live imaging microscopy (SCLIM), SAMs were dissected and visualized without fixation. GFP was excited with the 473-nm laser. The fluorescence emission spectra were separated with the custom-made dichroic mirror and filtered through a 490-545 bandpass filter. Images were acquired with an ImagEM EM-CCD camera (Hamamatsu Photonics). High-resolution images were constructed via deconvolution analysis performed with Volocity (PerkinElmer). A three-dimensional SCLIM image was constructed from 40 images taken at 0.1-µm vertical intervals using a theoretical point-spread function optimized for CSU10 confocal microscopes (Yokogawa Electric). *N. benthamiana* epidermal leaf cells were visualized with a confocal laser-scanning microscope (LSM 880; Zeiss, Tokyo, Japan) equipped with a 488-nm source and a 63× glycerol-immersion objective lens. For GFP fluorescence, images were captured at 500– 600 nm after excitation at 488 nm. After image acquisition, the images were processed using Zen software (Zeiss).

### Recombinant protein production and purification

The coding sequence of *Hd3a* and RFT1 were cloned into the pCold-GST vector^42^ and produced as a glutathione S-transferase (GST) fusion protein in *Escherichia coli* BL21 Rosetta (DE3) (Novagen). Cells were grown in LB medium or minimal medium containing 0.5 g l^™1^ of ^15^N-ammonium chloride. Recombinant GST fusion protein was purified with glutathione Sepharose 4B resin (Cytiva). After removal of the GST tag using GST-HRV 3C protease, Hd3a was purified by gel filtration chromatography using a Superdex75 column (Cytiva) with 50 mM potassium phosphate buffer (pH 6.8) containing 50 mM KCl and 1 mM dithiothreitol (DTT). To prepare Hd3a point mutants, plasmids were constructed by PCR using KOD plus DNA polymerase (TOYOBO) and the resulting recombinant proteins were purified essentially as described above.

The coding sequence of the 14-3-3 gene *GF14c*^*43*^ was cloned into the pCold-GST vector. GF14c was produced as a GST fusion protein in *E. coli* BL21 Rosetta (DE3) and purified with Glutathione Sepharose 4B resin and a Superdex 200 column (Cytiva), followed by removal of the GST tag.

The coding sequence of *OsFD1*^*2*^ was cloned into the pCold-GST vector. We could not purify the large amount of recombinant full-length protein. We therefore turned to a N-terminal-truncated version of OsFD1 (147–195) (lacking amino acids 1–146) with the S192E mutation, which was produced as a GST fusion protein in *E. coli* BL21 Rosetta (DE3) and purified as above.

### In vitro protein phase separation

Polyethylene glycol 8,000 (BioUltra grade, Sigma-Aldrich) was added to protein samples to a final concentration of 15% (w/v) in 25 mM potassium phosphate buffer (pH 6.8) containing 25 mM KCl and 0.5 mM DTT. The solutions were incubated for 24 h at 4 °C for protein phase separation. Phase separation was then assessed by measuring protein solution turbidity from 200 μl of protein solution (absorbance scanned between 330 and 800 nm) in 96-well plates (Stem, Tokyo) with a spectral scanning multimode reader (VarioSkan Flash 2.4, Thermo Fisher). Total protein concentration was 100 μM for all samples except for R121H mutant where 50 μM was used due to the difficulty in concentrating the protein solution. For phase-contrast image analyses, a Leica DMI300B microscope was used. The droplets of condensed Hd3a and RFT1 proteins were spherical and were hard to merge with each other even over a long lifetime, much like aging Maxwell glass.^44^.

### Nuclear magnetic resonance (NMR) experiments

All NMR spectra were acquired on an AVANCE III HD 800 spectrometer (Bruker) at 303 K, processed and analysed with TopSpin 4.11 software. ^1^H-^15^N HSQC experiments on 0.057 mM ^15^N-labelled Hd3a mutants were performed in 50 mM potassium phosphate buffer (pH 6.8) containing 50 mM KCl and 1 mM DTT.

### Protoplast transformation

Transformation of rice protoplasts was performed as described previously^2,28^. Five micrograms of *Hd3a* and *OsFD1* effector constructs and 0.5 µg of pUbqFluc reporter plasmid^2^ were transfected into 0.3–1.0 × 10^7^ protoplasts per ml by the PEG-mediated transformation method. After a 24-h incubation at 30 °C, the protoplast suspension was centrifuged and the cell pellet was frozen at ™80 °C for measurement of luciferase activity.

### Measurement of luciferase activity in protoplast lysates

Luciferase activity was measured as described previously^28^. The activities derived from NanoLuc (Nluc) and firefly luciferase (Fluc) in the lysates were measured separately and their ratio calculated. The Nano-Glo Luciferase Assay System (Promega) and Dual Luciferase Reporter Assay System (Promega) were used according to the manufacturer’s instructions for Nluc and Fluc activity, respectively. Transfected protoplasts were lysed with 25 µl of Passive lysis buffer (Promega). Nluc luminescence was measured with a TriStar LB941 microplate reader (Berthold Technologies) immediately after mixing of 10 µl of lysate and 10 µl of Nano-Glo Luciferase Assay Reagent in a 96-well plate. Fluc luminescence was measured immediately after mixing 2 µl of lysate and 10 µl of LARII Reagent in a 96-well microtiter plate.

### Data availability

The plasmids for transient expressions, plant transformations and protein purifications are available upon requests. Any other relevant data are available from the corresponding authors upon reasonable request.

## Figure legends

**Extended Data Fig. 1.**
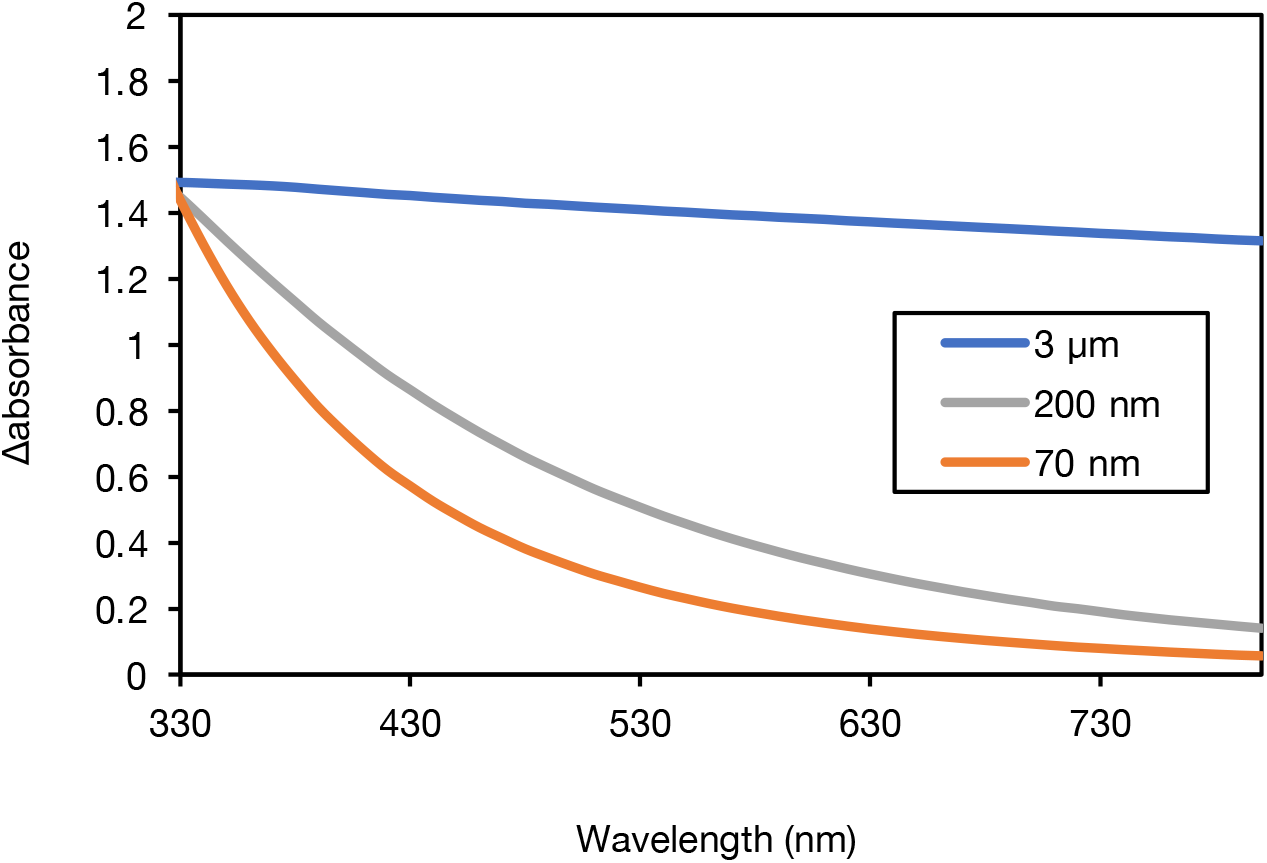
Changes in the absorbance of suspensions consisting of glass beads of different sizes. Absorbance between 330 and 800 nm of glass bead solutions with diameters of 3 µm, 200 nm and 70 nm.

**Extended Data Fig. 2.**
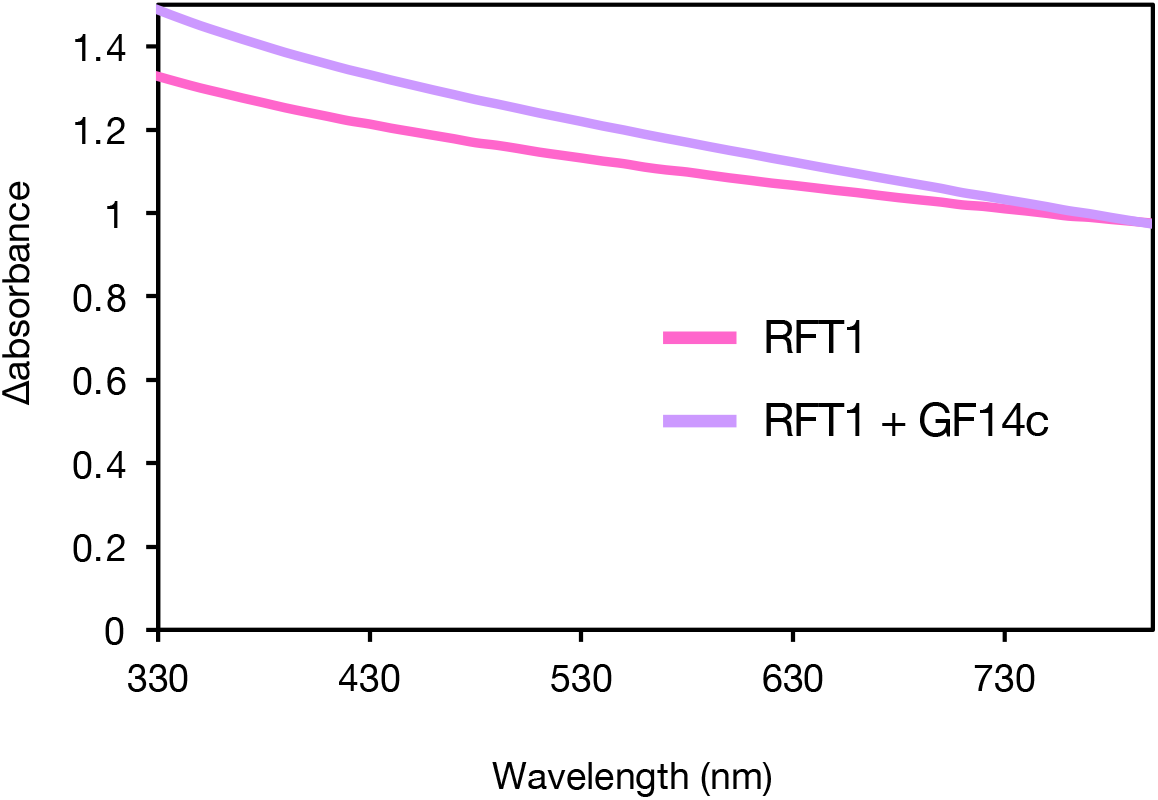
Absorbance of RFT1 with or without the 14-3-3 protein GF14c. Absorbance between 330 and 800 nm of recombinant RFT1 protein with (purple) or without (magenta) GF14c.

**Extended Data Fig. 3.**
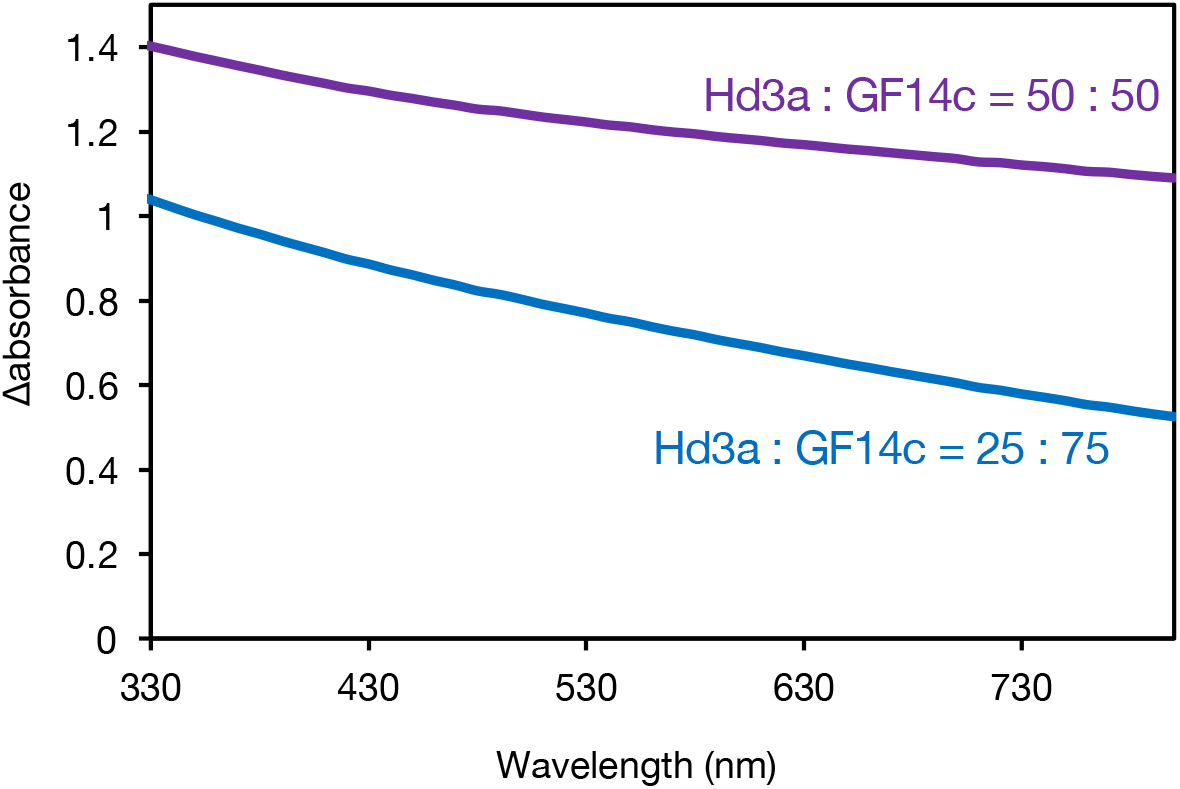
Ratio dependence of absorbance in Hd3a and GF14c. Absorbance between 330 and 800 nm of recombinant Hd3a with GF14c mixed in ratios of 50:50 (blue) or 25:75 (purple).

**Extended Data Fig. 4.**
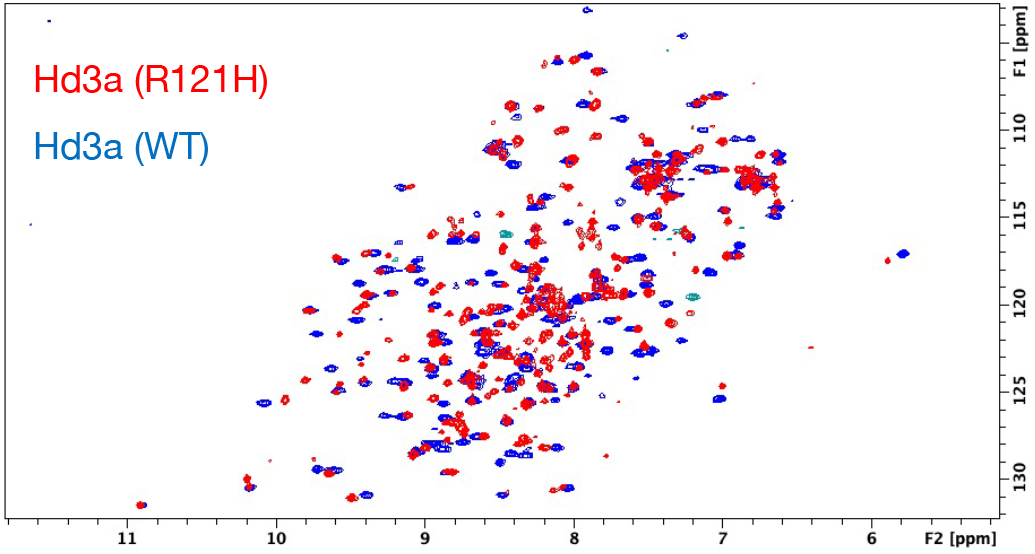
NMR spectra of Hd3a WT and R121H. ^1^H–^15^N heteronuclear single quantum coherence (HSQC) spectra of ^15^N-labelled recombinant Hd3a proteins for wild-type (WT, blue) and R121H mutant (red). p.p.m., parts per million.

## Acknowledgements

We thank Momoko Yoneyama for her technical support and Prof. Kentaro Shiraki for his comments on in vitro phase-separation experiments. This study was supported in part by MEXT/JSPS KAKENHI, Grant-in-Aid for Scientific Research (C), number 17K07609 to K.T., Grants-in-Aid for Scientific Research on Innovative Areas (numbers 16H06464 and 16H06466 to H.T., and 17H05836 and 19H04856 to C.K.), Grant-in-Aid for Scientific Research (B) (number 20H03191 to C.K.), Grant-in-Aid for Scientific Research (A) (number 16H02532 and 21H04728 to H.T.), a Core Research for Evolutionary Science and Technology (CREST) grant from the Japan Science and Technology Agency (JST) JPMJCR16O4 to H.T., Platform for Drug Discovery, Informatics, and Structural Life Science by MEXT, Platform Project for Supporting Drug Discovery and Life Science Research (Basis for Supporting Innovative Drug Discovery and Life Science Research, BINDS; number JP21am0101072) through the Japan Agency for Medical Research and Development, and the Collaborative Research Program of the Institute for Protein Research, Osaka University (CR-20-02).

## Author contributions

K.T., H.T. and C.K. designed the research, interpreted data and wrote the manuscript with the inputs from all authors.. K.T. performed transient expression analysis. S.A., M.E., Y.K. performed phase-separation assays. K.F. performed NMR measurements. T.K. and T.F. supervised protein analysis. M.T. performed microscopic observations of transient assays and plant transformation. J.N., A.F., M.F. and K.K. performed super-resolution imaging under the supervision of H.T. and A.N.. T.O. generated H2B-GFP transgenic rice.

## Competing Interests statement

The authors declare no competing financial interests.

## Supplementary materials

Extended Data Figs. 1–4

## Notes

### Competing Interest Statement

The authors have declared no competing interest.

## References

1. Andrés, F. & Coupland, G. The genetic basis of flowering responses to seasonal cues. Nature Reviews Genetics vol. 13 627–639 (2012).

2. Taoka, K.-I. et al. 14-3-3 proteins act as intracellular receptors for rice Hd3a florigen. Nature 476, (2011).

3. Collani, S., Neumann, M., Yant, L. & Schmid, M. FT Modulates Genome-Wide DNA-Binding of the bZIP Transcription Factor FD. Plant Physiol. 180, 367–380 (2019).

4. Li, C., Lin, H. & Dubcovsky, J. Factorial combinations of protein interactions generate a multiplicity of florigen activation complexes in wheat and barley. Plant J. 84, 70–82 (2015).

5. Chailakhyan, M. K. & Others. Hormonal theory of plant development. Hormonal theory of plant development. (1937).

6. Zeevaart, J. A. D. Florigen coming of age after 70 years. Plant Cell 18, 1783–1789 (2006).

7. Tamaki, S., Matsuo, S., Wong, H. L., Yokoi, S. & Shimamoto, K. Hd3a protein is a mobile flowering signal in rice. Science 316, 1033–1036 (2007).

8. Corbesier, L. et al. FT Protein Movement Contributes to Long-Distance Signaling in Floral Induction of Arabidopsis. Science 316, 1030–1033 (2007).

9. Kojima, S. et al. Hd3a, a Rice Ortholog of the Arabidopsis FT Gene, Promotes Transition to Flowering Downstream of Hd1 under Short-Day Conditions. Plant Cell Physiol. 43, 1096–1105 (2002).

10. Tamaki, S. et al. FT-like proteins induce transposon silencing in the shoot apex during floral induction in rice. Proc. Natl. Acad. Sci. U. S. A. 112, E901–E910 (2015).

11. Abe, M. et al. FD, a bZIP protein mediating signals from the floral pathway integrator FT at the shoot apex. Science 309, 1052–1056 (2005).

12. Wigge, P. A. et al. Integration of spatial and temporal information during floral induction in Arabidopsis. Science 309, 1056–1059 (2005).

13. Hell, S. W. & Wichmann, J. Breaking the diffraction resolution limit by stimulated emission: stimulated-emission-depletion fluorescence microscopy. Opt. Lett. 19, 780–782 (1994).

14. Klar, T. A., Jakobs, S., Dyba, M., Egner, A. & Hell, S. W. Fluorescence microscopy with diffraction resolution barrier broken by stimulated emission. Proc. Natl. Acad. Sci. U. S. A. 97, 8206–8210 (2000).

15. Kobayashi, Y., Kaya, H., Goto, K., Iwabuchi, M. & Araki, T. A pair of related genes with antagonistic roles in mediating flowering signals. Science 286, 1960–1962 (1999).

16. Kardailsky, I. et al. Activation tagging of the floral inducer FT. Science 286, 1962–1965 (1999).

17. Guivarc’H, A., Spena, A., Noin, M., Besnard, C. & Chriqui, D. The pleiotropic effects induced by therolC gene in transgenic plants are caused by expression restricted to protophloem and companion cells. Transgenic Res. 5, 3–11 (1996).

18. Searle, I. et al. The transcription factor FLC confers a flowering response to vernalization by repressing meristem competence and systemic signaling in Arabidopsis. Genes Dev. 20, 898–912 (2006).

19. Abiko, M., Maeda, H., Tamura, K., Hara-Nishimura, I. & Okamoto, T. Gene expression profiles in rice gametes and zygotes: identification of gamete-enriched genes and up-or down-regulated genes in zygotes after fertilization. J. Exp. Bot. 64, 1927–1940 (2013).

20. Kurokawa, K., Ishii, M., Suda, Y., Ichihara, A. & Nakano, A. Chapter 14 - Live Cell Visualization of Golgi Membrane Dynamics by Super-resolution Confocal Live Imaging Microscopy. in Methods in Cell Biology (eds. Perez, F. & Stephens, D. J.) vol. 118 235–242 (Academic Press, 2013).

21. Kurokawa, K. et al. Visualization of secretory cargo transport within the Golgi apparatus. J. Cell Biol. 218, 1602–1618 (2019).

22. Emenecker, R. J., Holehouse, A. S. & Strader, L. C. Emerging Roles for Phase Separation in Plants. Dev. Cell 55, 69–83 (2020).

23. Kim, J., Lee, H., Lee, H. G. & Seo, P. J. Get closer and make hotspots: liquid-liquid phase separation in plants. EMBO Rep. 22, e51656 (2021).

24. MacAlister, C. A. et al. Synchronization of the flowering transition by the tomato TERMINATING FLOWER gene. Nat. Genet. 44, 1393–1398 (2012).

25. Jung, J.-H. et al. A prion-like domain in ELF3 functions as a thermosensor in Arabidopsis. Nature 585, 256–260 (2020).

26. Fang, X. et al. Arabidopsis FLL2 promotes liquid-liquid phase separation of polyadenylation complexes. Nature 569, 265–269 (2019).

27. Huang, S., Zhu, S., Kumar, P. & MacMicking, J. D. A phase-separated nuclear GBPL circuit controls immunity in plants. Nature 594, 424–429 (2021).

28. Taoka, K.-I. et al. Novel assays to monitor gene expression and protein-protein interactions in rice using the bioluminescent protein, NanoLuc. Plant Biotechnol. 38, 89–99 (2021).

29. Cerise, M. et al. OsFD4 promotes the rice floral transition via florigen activation complex formation in the shoot apical meristem. New Phytol. 229, 429–443 (2021).

30. Serre, L., Vallée, B., Bureaud, N., Schoentgen, F. & Zelwer, C. Crystal structure of the phosphatidylethanolamine-binding protein from bovine brain: a novel structural class of phospholipid-binding proteins. Structure 6, 1255–1265 (1998).

31. Banfield, M. J., Barker, J. J., Perry, A. C. & Brady, R. L. Function from structure? The crystal structure of human phosphatidylethanolamine-binding protein suggests a role in membrane signal transduction. Structure 6, 1245–1254 (1998).

32. Hanzawa, Y., Money, T. & Bradley, D. A single amino acid converts a repressor to an activator of flowering. Proc. Natl. Acad. Sci. U. S. A. 102, 7748–7753 (2005).

33. Koornneef, M., Hanhart, C. J. & van der Veen, J. H. A genetic and physiological analysis of late flowering mutants in Arabidopsis thaliana. Mol. Gen. Genet. 229, 57–66 (1991).

34. Ahn, J. H. et al. A divergent external loop confers antagonistic activity on floral regulators FT and TFL1. EMBO J. 25, 605–614 (2006).

35. Ho, W. W. H. & Weigel, D. Structural Features Determining Flower-Promoting Activity of Arabidopsis FLOWERING LOCUS T. Plant Cell 26, 552–564 (2014).

36. Nakagawa, T. et al. Development of series of gateway binary vectors, pGWBs, for realizing efficient construction of fusion genes for plant transformation. J. Biosci. Bioeng. 104, 34–41 (2007).

37. Hellens, R. P., Edwards, E. A., Leyland, N. R., Bean, S. & Mullineaux, P. M. pGreen: a versatile and flexible binary Ti vector for Agrobacterium-mediated plant transformation. Plant Mol. Biol. 42, 819–832 (2000).

38. Nakamura, S. et al. Gateway binary vectors with the bialaphos resistance gene, bar, as a selection marker for plant transformation. Biosci. Biotechnol. Biochem. 74, 1315–1319 (2010).

39. Taoka, K.-I., Ham, B.-K., Xoconostle-Cázares, B., Rojas, M. R. & Lucas, W. J. Reciprocal phosphorylation and glycosylation recognition motifs control NCAPP1 interaction with pumpkin phloem proteins and their cell-to-cell movement. Plant Cell 19, 1866–1884 (2007).

40. Takeda, A. et al. Identification of a novel RNA silencing suppressor, NSs protein of Tomato spotted wilt virus. FEBS Lett. 532, 75–79 (2002).

41. Higo, A. et al. DNA methylation is reconfigured at the onset of reproduction in rice shoot apical meristem. Nat. Commun. 11, 4079 (2020).

42. Hayashi, K. & Kojima, C. pCold-GST vector: a novel cold-shock vector containing GST tag for soluble protein production. Protein Expr. Purif. 62, 120–127 (2008).

43. Purwestri, Y. A., Ogaki, Y., Tamaki, S., Tsuji, H. & Shimamoto, K. The 14-3-3 protein GF14c acts as a negative regulator of flowering in rice by interacting with the florigen Hd3a. Plant Cell Physiol. 50, (2009).

44. Jawerth, L. et al. Protein condensates as aging Maxwell fluids. Science 370, 1317–1323 (2020).

